# γδT Cells Are Required for CD8^+^ T Cell Response to Vaccinia Viral Infection

**DOI:** 10.1101/2021.04.01.438088

**Authors:** Rui Dai, Xiaopei Huang, Yiping Yang

## Abstract

Vaccinia virus (VV) is the most studied member of the poxvirus family, is responsible for the successful elimination of smallpox worldwide, and has been developed as a vaccine vehicle for infectious diseases and cancer immunotherapy. We have previously shown that the unique potency of VV in the activation of CD8^+^ T cell response is dependent on efficient activation of the innate immune system through Toll-like receptor (TLR)-dependent and -independent pathways. However, it remains incompletely defined what regulate CD8^+^ T cell response to VV infection. In this study, we showed that γδT cells play an important role in promoting CD8^+^ T cell response to VV infection. We found that γδT cells can directly present viral antigens in the context MHC-I for CD8^+^ T cell activation to VV *in vivo*, and we further demonstrated that cell-intrinsic MyD88 signaling in γδT cells is required for activation of γδT cells and CD8^+^ T cells. These results illustrate a critical role for γδT cells in the regulation of adaptive T cell response to viral infection and may shed light on the design of more effective vaccine strategies based on manipulation of γδT cells.

**Importance:** Targeting the immune systems has powerful potentials to treat many disorders, such as some cancers and viral infections. By understanding how the immune system responds to model infections, we can better determine strategies to manipulate our immune systems. Vaccinia virus is responsible for the worldwide elimination of smallpox and produces one of the longest immune responses known in humans. We know from previous findings that NK cells are required for initial immune response and CD8^+^ T cells are required for the elimination of the virus. How CD8^+^ T cells are activated in response to Vaccinia virus is not fully understood. This manuscript found that γδT cells activate CD8^+^ T cells in response to Vaccinia virus infection through MyD88 pathway

## Introduction

Vaccinia virus (VV), an enveloped double-stranded DNA virus, is a member of the *Orthoproxvirus* genus of the Poxviridae family. It has approximately a 200kb genome that encodes all the proteins required for cytoplasmic viral replication in host cells [1]. It is responsible for the worldwide elimination of smallpox, and as a result has been developed as recombinant vaccine vehicle for infectious diseases and cancer immunotherapy [2]. It is unique among viral agents to be able to elicit both potent and long-lasting immunity [3]. Though its natural route of infection is via the skin, many studies have noted that intraperitoneal, intravenous, and intramuscular modes of VV inoculation provides similar clinical efficacy in both mice and humans [4–6].

We have previously shown that the unique potency of intraperitoneal VV inoculation in the activation of CD8^+^ T cell responses is dependent on efficient activation of the innate immune system through Tolllike receptor (TLR)-dependent and -independent pathways [7, 8]. Specifically, we have demonstrated that intrinsic TLR2-MyD88 (myeloid differentiating factor 88) signaling in CD8^+^ T cells is critical for clonal expansion and long-lived memory formation [9]. In addition, TLR-independent production of type I interferons (IFNs) is also important for efficient CD8^+^ T cell responses [10, 11]. However, despite these advances, the mechanisms by conventional antigen-presenting cells are unable to fully explain the unique potency of VV in the activation of CD8^+^ T cell responses.

γδT cells are a unique population of lymphocytes that exert a strong influence on the immune system [12]. Previous studies have also shown that there are several subpopulations of γδT cells with distinct functions. However, a definitive system to categorize the different subpopulations has remained elusive. Initial proposals to define γδT cells based on their different TCR expression in both mice and humans have had to be revisited [13–18]. As a result, we assessed the effects of γδT cells as a whole for this study.

γδT cells act as a bridge between the innate and adaptive immune responses, with characteristics of both. They can exert direct cytotoxicity and enhance the adaptive immune responses [19–22]. Studies have found that γδT cells are important in the immune response against many mycobacterial, parasitic, and viral infections [23–33]. Similarly, previous studies have demonstrated that γδT cells express CD80 and CD86 at similar levels to professional antigen presenting cells and is able to promote CD8^+^ T cells activation [20, 21]. However, it remains largely unknown exactly how γδT cells promote adaptive immune responses.

In this study, we found that γδT cells play a critical role in promoting CD8^+^ T cell response to VV infection. We showed that activation of γδT cells by VV presented viral antigens in the context of MHC class I for CD8^+^ T cell activation *in vivo*. We further demonstrated that cell-intrinsic MyD88 signaling in γδT cells is required for γδT cell activation and CD8^+^ T cell responses. These results demonstrated a critical role for γδT cells in the regulation of CD8^+^ T cell response to viral infection and may shed light on the design of more effective vaccine strategies based on manipulation of γδT cells.

## Materials and Methods

### Animals

Eight-to ten-week-old C57BL/6, *δTCR*^−/−^, *OT-1*, and *β2m*^−/−^ mice were purchased from The Jackson Laboratory. *MyD88*^−/−^ on C57BL/6 background were kindly provided by Shizuo Akira (Osaka University, Osaka, Japan). All experiments involving the use of mice were done in accordance with protocols approved by the Animal Care and Use Committee at Duke University and the Ohio State University.

### Vaccinia virus

Western Reserve (WR) strain of VV was purchased from American Type Culture Collection (Manassas, VA). Recombinant VV-OVA was provided by Jonathan Yewdell at NIH. The viruses were grown in TK-143B cells and purified by centrifugation through a 35% sucrose cushion as previously described [34]. The titer was determined by plaque assay on TK-143B cells and subsequently stored at −80°C until use. For *in vivo* studies, 5×10^6^ pfu of live VV in 0.1mL Tris-Cl was injected into mice intraperitoneally, unless otherwise specified.

### DC culture

Femurs and tibiae of mice were harvest and bone marrow cells were flushed with DC medium (RPMI-1640 with 5% fetal bovine serum [FBS], 2mM L-glutamine, 10mM HEPES, 50μM β-mercaptoethanol, 100 IU/mL penicillin, and 100 IU/mL streptomycin), as previously described [34]. After lysis of red blood cells with ACK lysis buffer (Gibco Life Technologies, Waltham, MA), the bone marrow cells were culture in 6-well plates at density of 3×10^6^ cells/mL in 3mL DC medium in the presence of mouse granulocyte macrophage-colony stimulating factor (GM-CSF; 1000 U/mL; R&D Systems, Minneapolis, MN) and interleukin 4 (IL-4; 500 U/mL; R&D Systems). GM-CSF and IL-4 were replenished on day 2 and 4. On day 5, DCs were harvested, and CD11c^+^ DCs were transferred onto a new 24-well plate at a density of 0.85 × 10^6^ cells/mL in 2mL DC media.

### Isolation of γδT cells

Splenocytes were harvested from C57BL/6 mice 2 days after peritoneal inoculation with VV. γδT cells were isolated form harvested splenocytes with pan-T cell microbeads, followed by anti-γδTCR microbeads (Miltenyi Biotec, Auburn, CA). The isolated γδT cells were assessed via flow cytometry for confirmation.

### CD8^+^ T cell proliferation assay

CD8^+^ T cells were isolated from splenocytes of OT-I mice on C57BL/6 background using anti-CD8a microbeads (Miltenyi Biotec), and then fluorescently labeled with carboxyfluorescein succinimidyl ester (CFSE). Labeled CD8^+^ T cells and OVA-I peptide were then cocultured with matured DCs or VV-activated γδT cells at 1:1 ratio in 96 well plates. The cells were incubated at 37°C for 72 hours, and then assessed via flow cytometry.

### Adoptive transfer of γδT cells

Naïve γδT cells were isolated from pooled spleens and lymph nodes of wild-type or *MyD88*^−/−^ mice on C57BL/6 background, with pan-T cell microbeads, followed by anti-γδTCR microbeads (Miltenyi Biotec). The isolated cells were confirmed via flow cytometry for confirmation and suspended in phosphate buffered saline (PBS). The cells were then injected intravenously via the tail vein into *δTCR*^−/−^ or *MyD88*^−/−^ mice on C57BL/6 background at 1×10^6^ cells/mouse, unless otherwise specified.

### Antibodies and flow cytometry analysis

The list of used antibodies is provided in Table 1. Cells were suspended in PBS buffer with 2% heat-inactivated FBS and 0.1% sodium azide. After staining, cells were washed twice, and analyzed with FACSCanto flow cytometer (BD Biosciences) using FlowJo software (BD Biosciences)

**Table 1 –.** Antibodies

| **Antibody** | **Fluorophore** | **Company** | **Catalog Number** |
| --- | --- | --- | --- |
| CD3e | APC | BD | 553066 |
| CD4 | APC | BD | 553051 |
| IFN-γ | APC | BioLegend | 505810 |
| MHC-I H2D | APC | eBioscience | 17-5998-30 |
| CD4 | FITC | BD | 553047 |
| CD8a | FITC | BD | 553031 |
| TCR$\gamma\delta$ | FITC | eBioscience | 11-9959-42 |
| CD80 (B7.1) | PE | BioLegend | 104709 |
| CD86 (B7.2) | PE | BD | 553692 |
| H-2k(b) TSYKFESV | PE | NIH | 7716 |
| CD4 | PE | BD | 553730 |
| CD8a | PE | BD | 553032 |
| TCR$\gamma\delta$ | PE | BioLegend | 118107 |
| H2Kb SIINFEKL | PE | BioLegend | 141603 |
| CD3e | PE-Cy5 | BD | 553065 |
| CD4 | PE-Cy5 | BD | 553050 |
| CD8a | PE-Cy5 | BD | 553034 |
| TCR$\gamma\delta$ | PE-Cy7 | eBioscience | 25-5711-80 |
| CD4 | PE-Cy7 | BioLegend | 100421 |

### Intracellular cytokine staining

Splenocytes were re-stimulated specifically for CD8^+^ T cells with 2 μg/mL B8R peptide (TSYKFESV, BD Pharmingen) with 5 μg/mL Brefeldin A (Invitrogen) for 5 hours at 2 μg/mL at 37°C. Splenocytes or mesenteric lymph node cells were stimulated specifically for γδT cells with 50 ng/mL Ionomycin, 100 ng/mL PMA, and 5 μg/mL Brefeldin A for 3 hours at 37°C. After staining with cell surface markers, the cells were fixed and permeabilized with Cytoperm/Cytofix solution (BD Biosciences) for 20 minutes and incubated with anti-IFN-γ antibodies for 30 minutes. The cells were washed twice with Permeabilization buffer (BD Biosciences) and analyzed with a FACSCanto flow cytometer using FlowJo software (BD Biosciences)

### MHC/peptide tetramer

The VV-specific epitope B8R_20-27_, TSYKFESV, is a synthetic peptide based on modified vaccinia virus Ankara (MVA) sequence [35]. Peptide MHC I tetramers consisting of B8R_20-27_/K^b^ conjugated to allophycocyanin were obtained from the NIH Tetramer Core Facility (Emory University, Atlanta, GA, USA). Cells were stained with the tetramer for 30 minutes at room temperature in the dark together with surface staining and subsequently analyzed by flow cytometry.

### Statistical analysis

Results are expressed as mean ± SEM. Differences between groups were examined for statistical significance using Kolmogorov-Smirnov test, Mann-Whitney test, or unpaired t-test with Welch’s correction. *P*-values less than 0.05 are considered to be significant.

## Results

### γδT cells are required for CD8^+^ T cell response to VV

To address whether γδT cells play a role in regulating CD8^+^ T cell responses, we first examined the activation status of γδT cells in response to VV infection in vivo. C57BL/6 mice were injected with VV intraperitoneally, and at different time points after infection, γδT cells were examined for IFN-γ production. We found that in both spleen (Fig. 1A) and peritoneal cavity (Fig. 1B), IFN-γ^+^ γδT cell count reached its peak around day 4 following VV infection, with subsequent decline in the days following. This is in contrast to VV-specific CD8^+^ T cell response in that IFN-γ^+^ CD8^+^ T cell count reached its peak around day 7 (Fig. 1). These results indicated that the activation of γδT cells peaked prior to that of CD8^+^ T cells. We found that naïve γδT cells secreted IFN-γ after stimulation, therefore we subsequently always comparatively determined IFN-γ gating within each experiment between inoculated versus naïve mice (Fig. 1C, D).

**FIGURE 1.**
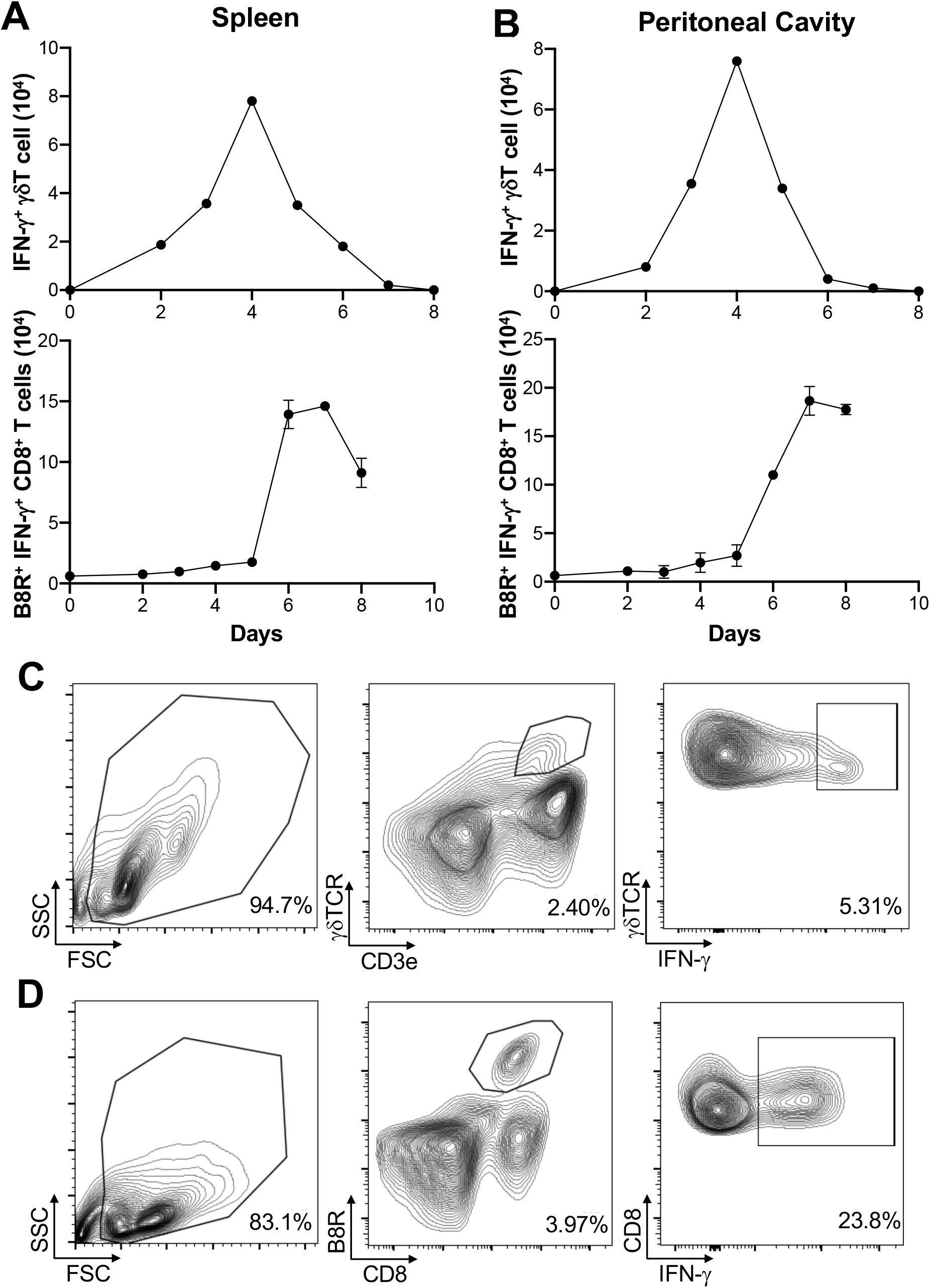
Activation of γδT cells in response to VV infection. 5 × 10^6^ pfu of VV were injected into wildtype C57BL/6 mice intraperitoneally. 2, 3, 4, 5, 6, 7 and 8 days after infection, CD8^+^ T cells from spleen and peritoneal cavity were assayed for IFN-γ production by intracellular staining by FACS, and IFN-γ^+^ γδT cells and CD8^+^ T cells from the spleen (**A**) spleen and peritoneal cavity (**B**) are shown. IFN-γ^+^ γδT cells are first gated on CD3e^+^ and γδTCR^+^, then assessed on IFN-γ expression. VV-specific B8R^+^ IFN-γ^+^ CD8^+^ T cells were first gated on B8R^+^ CD8^+^ T cells, and then assessed for IFN-γ expression. The mean of each time point is plotted. Representative of 2 independent studies, each with 3 biological replicates. Gating strategy used to generate the data are represented by (**C**) and (**D**). The gating for IFN-γ^+^ γδT cells is determined against IFN-γ^+^ γδT cells from control naïve mice for each experiment. Mann-Whitney test, *P* < 0.05.

We next determined if γδT cells play a role in CD8^+^ T cell response to VV. We inoculated wild-type (WT) and *δTCR*^−/−^ C57BL/6 mice intraperitoneally with VV and assessed for VV-specific B8R^+^ CD8^+^ T cell activation 7 days post-inoculation. B8R is a VV epitope that is recognized by VV-specific CD8^+^ T cells; B8R^+^ CD8^+^ T cells are specifically activated by VV. We found that there is a significant decrease in VV-specific B8R^+^ (Fig. 2C, E) and functional IFN-γ^+^ (Fig. 2D, F) CD8^+^ T cells in *δTCR*^−/−^ mice that lack γδT cells, compared to that of WT mice (*P* < 0.005). We subsequently found that this defect can be rescued with adoptively transferred WT γδT cells. VV inoculation of *δTCR*^−/−^ mice with adoptive transfer of WT γδT cells had significantly greater VV-specific B8R^+^ and IFN-γ^+^ CD8^+^ T cells, compared to that of *δTCR*^−/−^ mice with VV inoculation alone (*P* < 0.005). VV-specific B8R^+^ and IFN-γ^+^ CD8^+^ T cell response in *δTCR*^−/−^ with adoptive transfer of WT γδT cells following VV inoculation also approximated the same response as WT mice with VV inoculation alone (*P* not significant). This suggested that γδT cells play a critical role in CD8^+^ T cell activation following VV infection.

**FIGURE 2.**
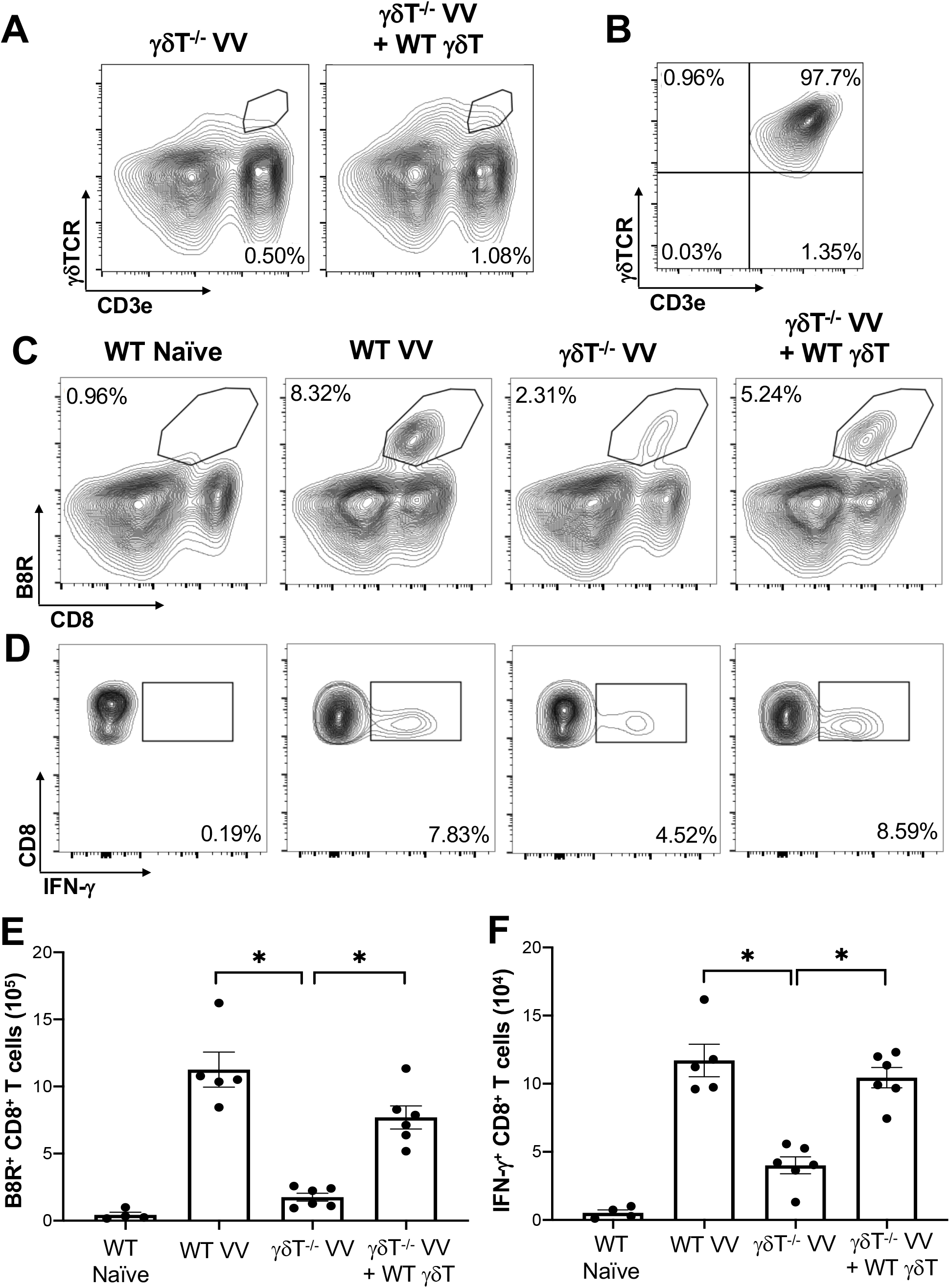
γδT cells is required for CD8^+^ T cell response to VV. 5 × 10^6^ pfu of VV were injected intraperitoneally into wild-type C57BL/6 (WT) or *δTCR*^−/−^ mice (γδT^−/−^). Concurrently, a different population of *δTCR*^−/−^ mice were also adoptively transferred with 1 × 10^6^ WT γδT cells (γδT^−/−^ VV + WT γδT), followed by VV inoculation. (**A**) 3 days post-adoptive transfer and VV inoculation, there is a detectable population of γδT cells in the mesenteric lymph nodes. (**B**) Representative plot of purified γδT cells used for adoptive transfer. 7 days post-inoculation, the spleens were harvested and stained for B8R^+^ CD8^+^ T cells by tetramer and IFN-γ^+^ CD8^+^ T cells by intracellular staining. (**C**) Representative FACS plots first gated on CD8^+^ CD4^−^ T cells and then plotted against B8R^+^ and CD8^+^ T cells for B8R^+^ CD8^+^ T cells. (**D**) Representative FACS plots first gated on CD8^+^ CD4^−^ T cells and then assessed for IFN-γ expression. (**E, F**) Quantification of FACS plots. Values are mean ± SEM, representative of 3 independent studies, each with at least 3 biological replicates. Kolmogorov-Smirnov nonparametric t-test, \**P* < 0.005.

### VV activates γδT cells to present MHC-I peptide and upregulate CD80 and CD86

To determine how γδT cells promote the activation CD8^+^ T cells to VV infection, we explored whether γδT cells contributed to signals that are required to activate CD8^+^ T cells: 1) direct presentation of VV-specific peptide on MHC-I, 2) co-stimulation with CD80 and CD86 ligands [36–38]. To assess peptide presentation on MHC-I, we inoculated WT mice with VV or VV encoded with OVA (VV-OVA). We then assessed γδT cells for expression of H2K^b^ specific for SIINFEKL peptide on MHC-I. We found that there is an increase in H2K^b^ SIINFEKL^+^ γδT cells in mice inoculated with VV-OVA, compared to that of naïve or mice inoculated with VV (Fig. 3A).

**FIGURE 3.**
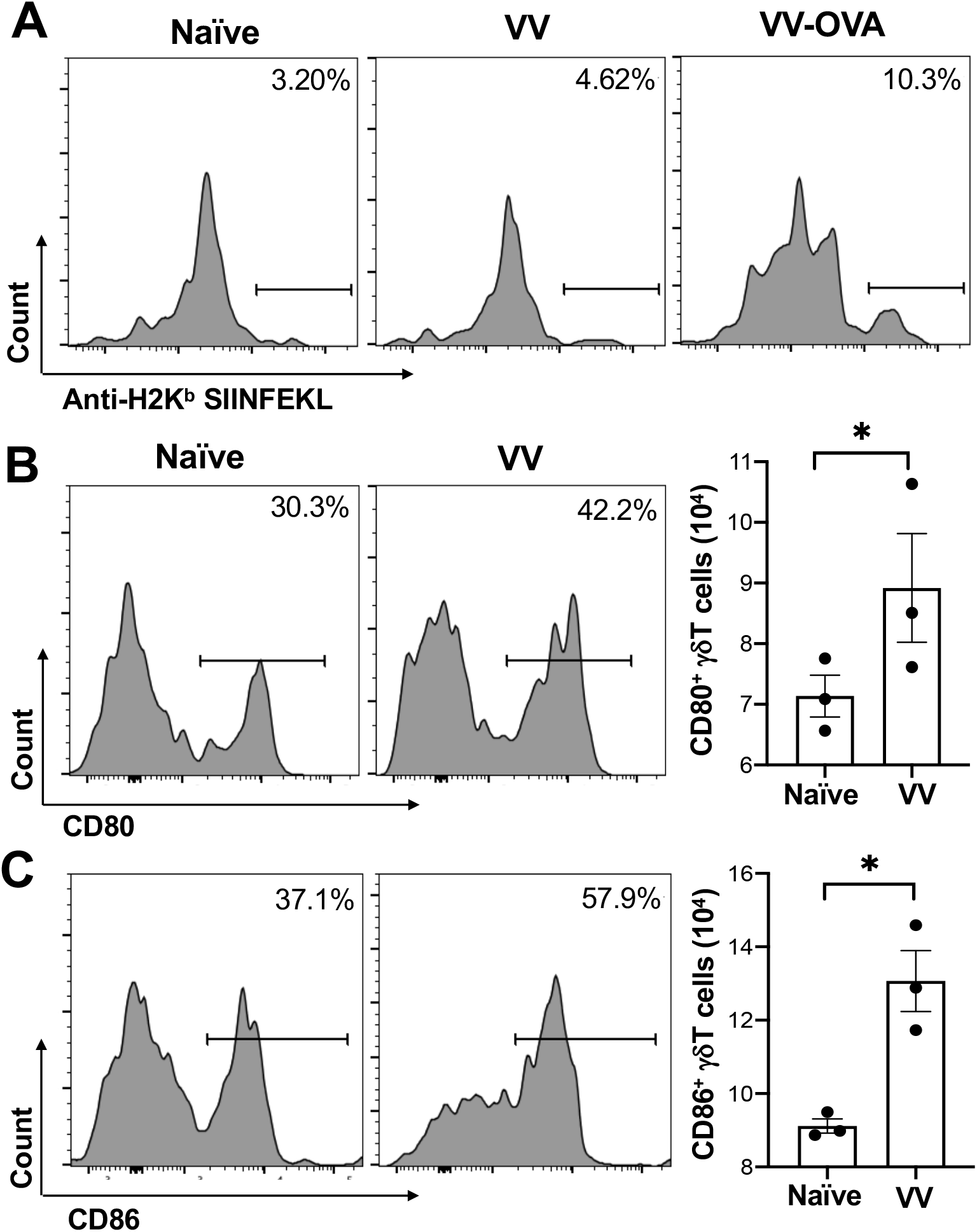
VV activates γδT cells to present peptide on MHC-I and upregulate CD80/CD86 expression. C57BL/6 mice were inoculated intraperitoneally with 5 × 10^6^ pfu of VV or VV encoding OVA (VV-OVA). (**A**) Representative FACS plots first gated on CD3e^+^ and γδTCR^+^ T cells, and then assessed for SIINFEKL peptide expression on mouse MHC-I H2K^b^ via anti-H2K^b^ SIINFEKL in spleens harvested 2 days post-infection. (**B, C**) 4 days post-inoculation, splenocytes were extracted and stained for CD3e^+^ and γδTCR^+^ T cells. The gated cells were then assessed for (**B**) CD80 and (**C**) CD86 expression. Values are mean ± SEM, representative of 3 independent studies, each with at least 3 biological replicates. Kolmogorov-Smirnov nonparametric t-test, \**P* < 0.05.

We also found that following VV infection, there is an increase in CD80 and CD86 expression on the surface of γδT cells by flow cytometry. CD86 is expressed first as the initial co-stimulatory ligand, and CD80 is expressed after antigen-presenting-cell activation [39]. We found that 4 days post-inoculation, there is a significant increase in CD86 (Fig. 3C; *P* < 0.01), and a corresponding increase in CD80 (Fig. 3B; *P* < 0.05). These results suggests that γδT cells could provide the necessary signals for CD8^+^ T cell activation after VV infection.

### γδT cells also directly activate CD8^+^ T cells via MHC-I

We next explored whether γδT cells directly activates CD8^+^ T cell following VV infection *in vivo*. To determine if VV can activate γδT cells to act as antigen presenting cells, we isolated γδT cells from WT C57BL/6 mice that had been inoculated intraperitoneally with VV 48 hours prior. CD8^+^ T cells were obtained from OT-I mice on C57BL/6 background and pulsed with CFSE. The CFSE-labeled CD8^+^ T cells were then co-cultured with the VV-activated γδT cells and OVA-I peptide. 72 hours after co-incubation, CD8^+^ T cell proliferation was assayed by CFSE dilution. As a control, the CD8^+^ T cells were also incubated with matured DCs and OVA-I peptide. We found that CD8^+^ T cells proliferated at a similar magnitude when cocultured with VV-activated γδT cells as with matured DCs (Fig. 4A). The same proliferation is not seen if CD8^+^ T cells are incubated with γδT cells or OT-I peptide alone. This suggests that VV is able to activate γδT cells to become antigen presenting cells.

**FIGURE 4.**
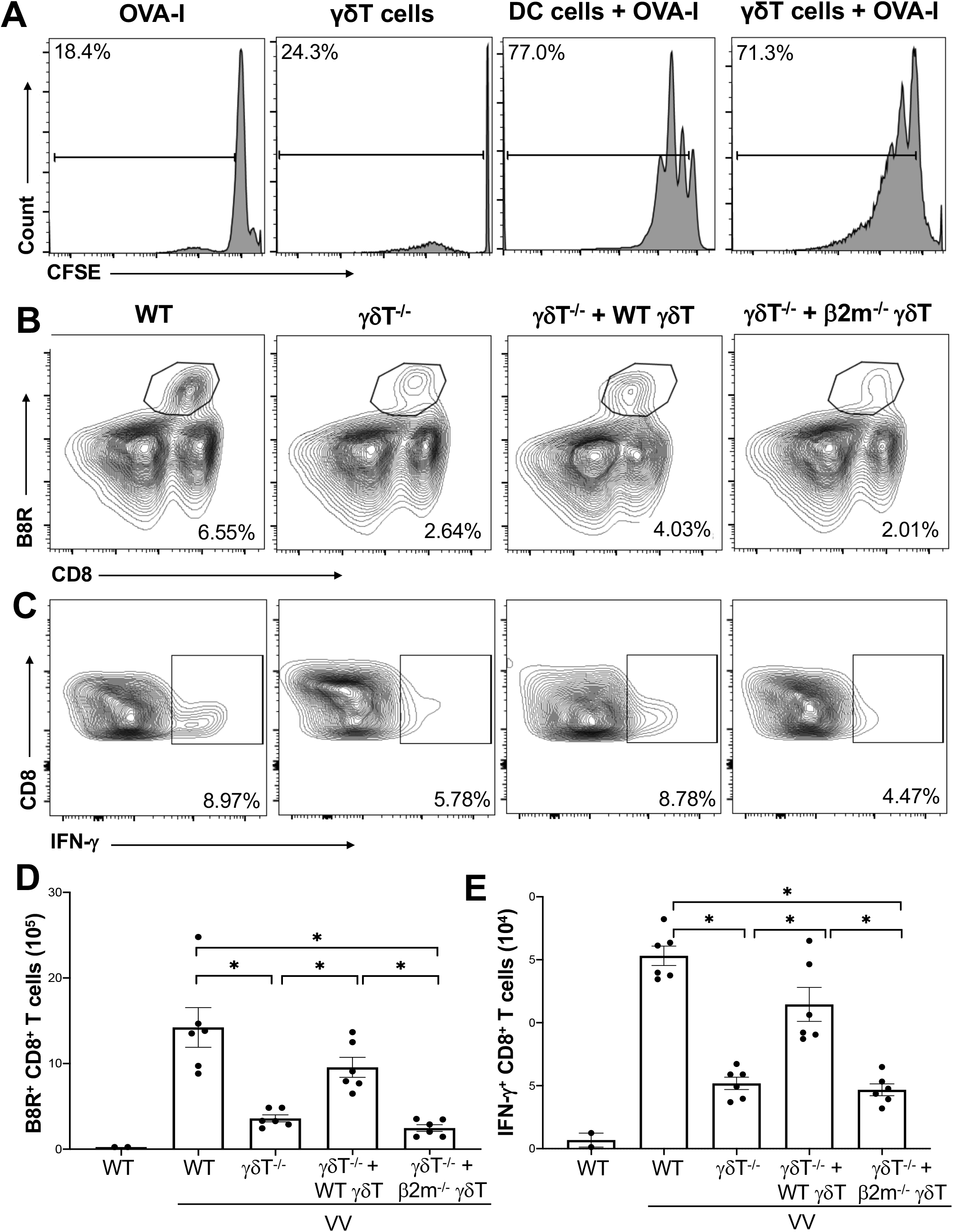
γδT cells directly activate CD8^+^ T cells via MHC-I. CD8^+^ T cells were obtained from splenocytes of OT-I^+^ mice via magnetic-activated cell sorting. (**A**) The CD8^+^ T cells were labeled with Carboxyfluorescein diacetate succinimidyl ester (CSFE) and measured for proliferation. The labeled cells were co-cultured *in vitro* with OVA-I peptide plus LPS-matured dendritic cells or VV-activated γδT cells for 3 days, at a ratio of 1:1. Cells were harvested and stained for CD8 and CD3e. Control CD8^+^ T cells is incubated with OT-I peptide or γδT cells alone. They were subsequently assessed via flow cytometry for CSFE. (**B**, **C**) 5 × 10^6^ pfu of VV was inoculated intraperitoneally into WT or *δTCR*^−/−^ mice, with and without adoptive transfer of 1 x10^6^ cells of WT or *β2m* γδT cells. 7 days post-inoculation, splenocytes were stained for CD8^+^ CD4^−^ lymphocytes, and assessed for B8R^+^ and IFN-γ^+^ CD8^+^ T cells. (**B**) Representative FACS plots first gated on gated on CD8^+^ CD4^−^ T cells and then plotted against B8R^+^ and CD8^+^ T cells for B8R^+^ CD8^+^ T cells. (**C**) Representative FACS plots first gated on CD8^+^ CD4^−^ T cells and then assessed for IFN-γ expression. **(D, E)** Quantification of FACS plots. Values are mean ± SEM, representative of 3 independent studies. Kolmogorov-Smirnov nonparametric t-test, \**P* < 0.005.

We then determined if γδT cells functionally acts as professional APCs via MHC-I by assessing if CD8^+^ T cell activation could be rescued via adoptive transfer of *β2m*^−/−^ γδT cells. β2m is a necessary component of MHC-I. We found that adoptive transfer of WT γδT cells into *δTCR*^−/−^ followed by VV infection significantly increased percentages and cell count of B8R^+^ and IFN-γ^+^ CD8^+^ T cells compared to *δTCR*^−/−^ with VV infection alone (Fig. 4B-E; *P* < 0.001). However, *β2m* γδT cells into *δTCR*^−/−^ followed by VV infection resulted in similar percentages and cell count of B8R^+^ and IFN-γ^+^ CD8^+^ T cells as *δTCR*^−/−^ with VV infection alone (Fig. 4B-E; *P* < 0.001). This suggests that γδT cells also activate CD8^+^ T cell via presentation of epitope on MHC-I for CD8^+^ T cell recognition.

### MyD88 signaling in γδT cells promotes CD8^+^ T cell response to VV

To determine how VV activates γδT cells for antigen presentation, we screened mice defective for innate signaling pathways and assessed for γδT cell proliferation and IFN-γ^+^ γδT cell functional activation. VV was inoculated in WT, *IFN-αβR*^−/−^, *IFN-γR*^−/−^, *TNF-αR*^−/−^, and *MyD88*^−/−^ mice and splenic cells were assessed via flow cytometry 4 days afterwards. We found that there was a significant deficit in γδT cell proliferation and IFN-γsecretion in *MyD88*^−/−^ mice (Fig. 5), but not in mice with other deficient signaling pathways. This suggests that MyD88 signaling is required for VV activation of γδT cells.

**FIGURE 5.**
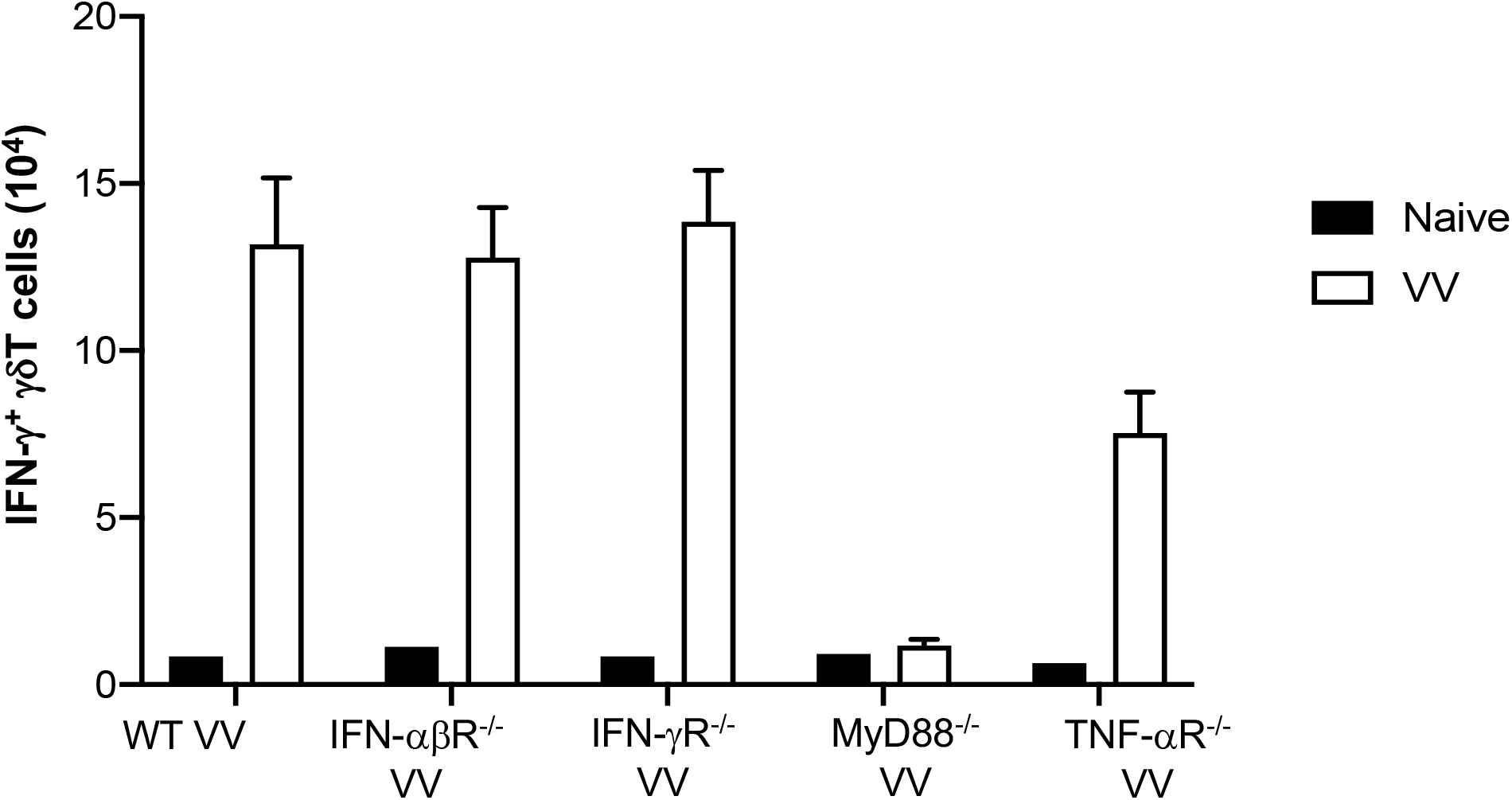
*In vivo* screen for signaling pathways required for γδT cell activation following VV infection. Wild-type (WT), IFN-αβ receptor deficient (IFN-αβR^−/−^), IFN-γ receptor deficient (IFN-γR^−/−^), MyD88 deficient (MyD88^−/−^), and TNF-α receptor deficient (TNF-αR^−/−^) mice were inoculated with VV intraperitoneally and splenocytes were harvested 4 days after inoculation. Harvested cells were gated for CD3e^+^ γδTCR^+^ T cells and subsequently assessed IFN-γ positivity.

We next addressed if intrinsic MyD88 activation in γδT cells is sufficient for VV activation, opposed to signaling from other cells, we adoptively transferred WT γδT cells into *MyD88*^−/−^ mice and assessed for γδT cell activation. We found that adoptive transfer of WT γδT cells into *MyD88*^−/−^ mice is sufficient to rescue γδT cell proliferation and IFN-γ^+^ secretion (Fig. 6; *P* < 0.001), suggesting intrinsic MyD88 signaling is required for activation of γδT cells by VV.

**FIGURE 6.**
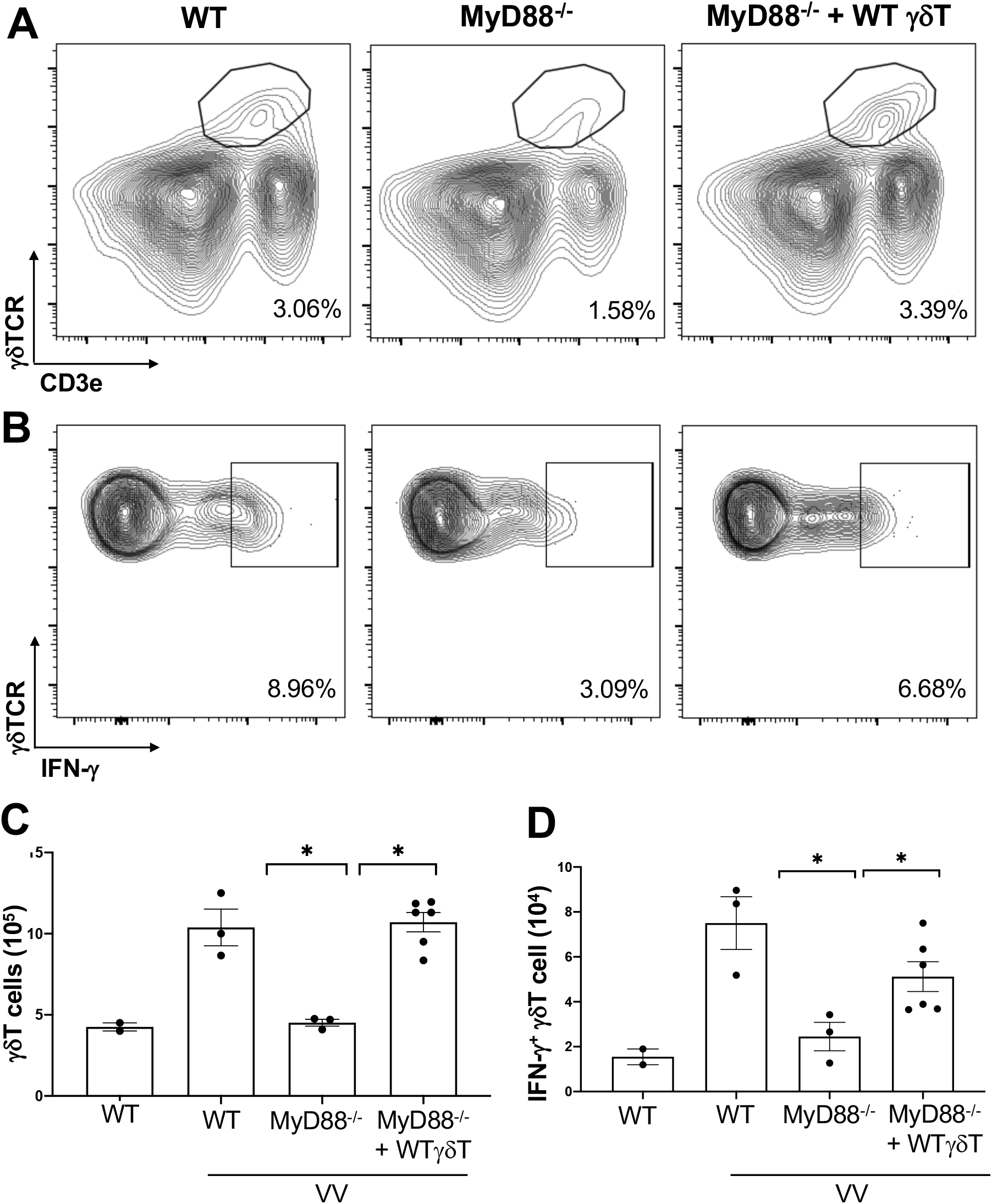
Intrinsic MyD88 signaling is required for activation of γδT cells by VV. 5 × 10^6^ pfu of VV was inoculated into WT and *MyD88* mice with and without adoptive transfer of 1 × 10^6^ cells of WT γδT cells. Mice splenocytes were harvested 4 days after inoculation and assessed via flow cytometry. (**A**) Representative FACS plots of live cells plotted for CD3e^+^ γδTCR^+^ splenocytes, 4 days after VV inoculation. (**B**) Representative FACS plots first gated on CD3e^+^ γδTCR^+^ splenocytes, and then assessed for IFN-γ positivity. The gating for IFN-γ is determined against γδT cells from control naïve mice for each experiment. (**C**) Summary graph of a representative experiment for positive CD3e^+^ γδTCR^+^ T cells in mice, 4 days post-VV inoculation. (**D**) Summary graph of a representative experiment for IFN-γ^+^ CD3e^+^ γδTCR^+^ T cells in mice, 4 days post-VV inoculation. Representative of 3 independent experiments. Kolmogorov-Smirnov nonparametric t-test, \**P* < 0.001,

We then investigated if MyD88 signaling in γδT cells is needed for subsequent CD8^+^ T cell activation. We adoptively transferred WT or *MyD88*^−/−^ γδT cells into *δTCR*^−/−^ mice and found that there was similar γδT cell reconstitution (Fig 7A), but that was a significant decrease in VV-specific B8R^+^ and IFN-γ^+^ CD8^+^ T cell proliferation in *δTCR*^−/−^ adoptively transferred with *MyD88*^−/−^ γδT cells when compared to that of WT γδT cells (Fig. 7B-E; *P* < 0.005).

**FIGURE 7.**
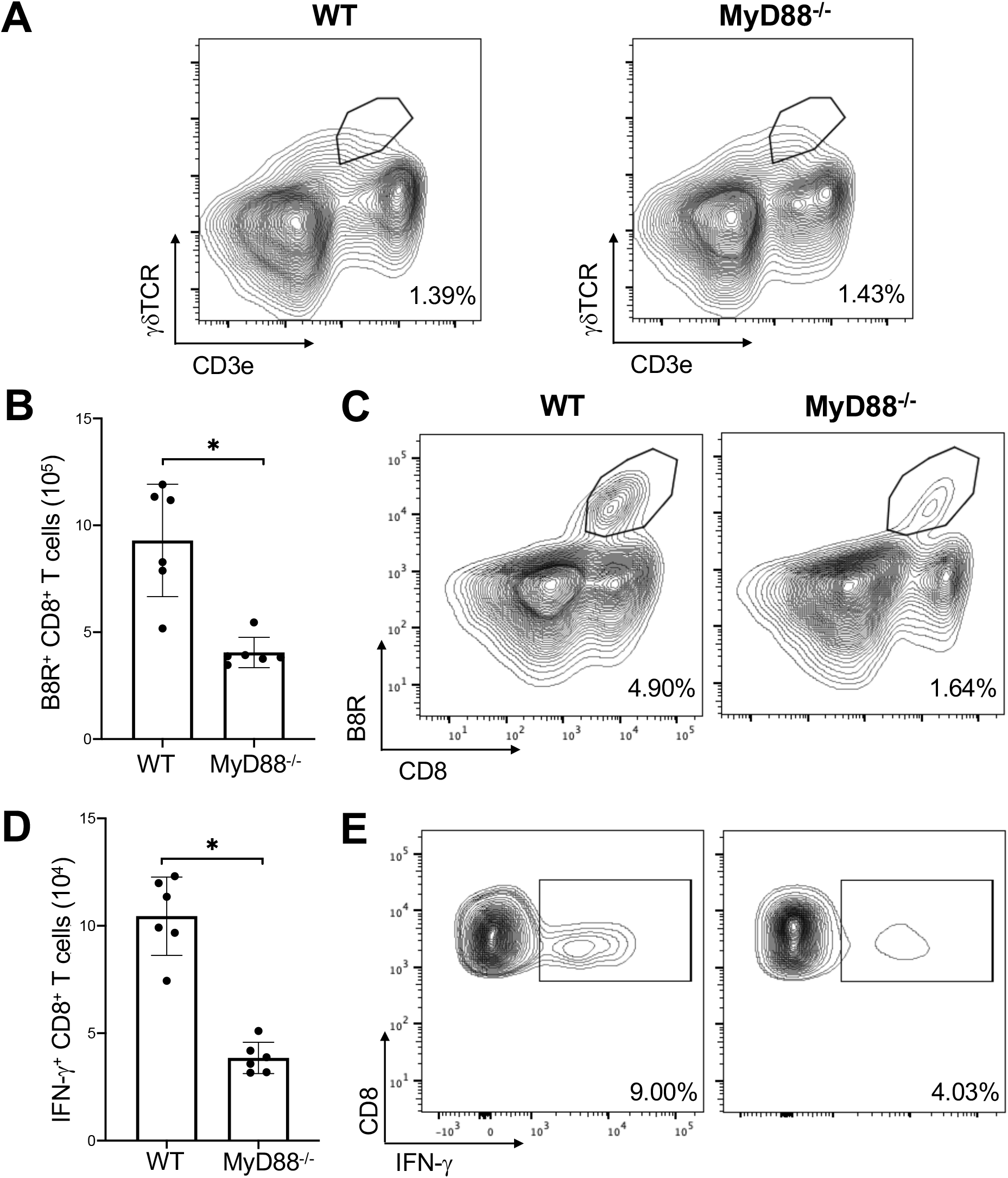
MyD88 is required for CD8^+^ T cell activation by γδT cells. *δTCR*^−/−^ mice were adoptively transferred with WT or *MyD88*^−/−^ γδT cells and inoculated with VV. (**A**) 3 days after inoculation, there is no significant difference between *δTCR*^−/−^ mice adoptively transferred with WT vs *MyD88*^−/−^ γδT cells. 7 days after inoculation, splenocytes were obtained and stained for CD8^+^ CD4^−^ lymphocytes and plotted via flow cytometry for (**B**) B8R^+^ CD8^+^ T cells or (**C**) IFN-γ^+^ CD8^+^ T cells. (**D**) Representative FACS plots of splenocytes first gated on CD8^+^ CD4^−^ lymphocytes, and then plotted for CD8^+^ B8R^+^ T cells. (**E**) Representative FACS plots of splenocytes first gated on CD8^+^ CD4^−^ lymphocytes, and then assessed for IFN-γ positivity. Unpaired student T-test, \**P* < 0.005. Each panel is representative of 3 independent studies.

## Discussion

Here we showed that VV can activate γδT cells via the MyD88 signaling pathway. We further showed that VV-activated γδT cells can present antigens to activate and induce VV-specific CD8^+^ T cell response. Our results further demonstrate MyD88 has a critical role in VV activation of γδT cells to promote specific CD8^+^ T cell response.

γδT cells represents approximately 0.7% of the peripheral blood and play an important role in the integration of the innate and adaptive immune system [19]. Previous studies have shown that although activation of NK cells is critical for the initial control of VV infection [7, 8], efficient activation of CD8^+^ T cell response is required for the eradication of VV infection [9]. What promotes the activation of CD8^+^ T cell response to VV infection remains incompletely defined. Understanding how γδT cells activate CD8^+^ T cells will better elucidate the mechanisms that govern CD8^+^ T cell activation and how to better employ them for future strategies in vaccination or immunotherapy.

γδT cells play an active role in the control of parasitic, bacterial, and viral infections, such as malaria, Listeria monocytogenes, Salmonella, EBV, and HSV [23, 25–28, 40–42]. Unlike other innate immune cells, γδT cells require activation by various antigens prior to exhibiting cytotoxic characteristics [43]. Currently, most strategies that target γδT cells employ phosphoantigens to activate γδT cells. However, recent evidence suggests that phosphoantigen activation is nonspecific, and induce both inflammatory and antiinflammatory functions in the targeted cells [44–47]. This may be due to the numerous subpopulations of γδT cells that exert balancing functions against each other [48–50]. The goal is therefore to investigate a method that would preferentially activate one subpopulation of γδT cells, however categorization of γδT cells remain controversial [14, 17]. Recent studies have shown that even γδT cells with the same γ and δ TCR chains appear to have distinctly different functions [51–54]. Given the long-lasting immunity that VV produce in clinical evaluations, understanding how VV activates γδT cells could provide insights into strategies to shifting γδT cells towards cytotoxicity immunity overall.

In this study, we demonstrate that γδT cells is required for the full activation of CD8^+^ T cells after VV infection. *δTCR*^−/−^ mice have deficient VV-specific CD8^+^ T cell proliferation and functional response. We find that deficiency of γδT cells results in over 3-fold decrease in CD8^+^ T cell response. Given previous studies that demonstrate γδT cells’ influence on the immune system, it is possible that γδT cells may be directly responsible for CD8^+^ T cell activation following VV infection [20, 21, 55]. To investigate this possibility, we investigated whether VV upregulates the 3 conventional signals of CD8^+^ T cell activation in γδT cells. We found that following infection with VV-OVA, SIINFEKL peptide is present on MHC-I on the surface of γδT cells. We also found that there is a significant decrease in CD8^+^ T cell activation in *δTCR*^−/−^ mice adoptively transferred with deficient MHC-I γδT cells, compared to that transferred with wild-type γδT cells. Similarly, there is a significant increase in expression of CD86, IFN-α, and IL-1 in γδT cells following VV infection for signals 2 and 3 that are required for CD8^+^ T cell activation. This suggests that γδT cells can provide all 3 signals necessary for CD8^+^ T cell activation, and that γδT cells directly influence CD8^+^ T cell activation following VV infection.

To determine how VV activates γδT cells for antigen presentation, we determined γδT cell activity in mice with deficiencies in IFN-αβ, IFN-γ, TNF-α, and MyD88 signaling. We found that only mice with deficient MyD88 signaling presented with impaired γδT cell response to VV infection. Subsequently, we investigated whether the γδT cell deficiency see in *MyD88* mice are due to signaling from other cells that require MyD88 signaling or MyD88 in γδT cells alone. We found that *MyD88* mice that were adoptively transferred with WT γδT cells exhibited normalized γδT cells response to VV. This indicates that VV activates γδT cells via MyD88-associated PRRs. Additionally, when *MyD88* γδT cells are adoptively transferred to *δTTCR*^−/−^ mice, there is a deficient CD8^+^ T cell response. This means that MyD88 activation on γδT cells is required for CD8^+^ T cell response.

In conclusion, our study reveals that VV activates γδT cells via MyD88 signaling pathway, which leads to direct antigen presentation to CD8^+^ T cells. *In vivo*, this bridge between the innate and adaptive immune pathways plays a critical role in the activation of CD8^+^ T cell response to VV. Furthermore, we demonstrate that cell-intrinsic MyD88 signaling in γδT cells is required for activation of CD8^+^ T cells. These results demonstrate a critical role for γδT cells in the regulation of adaptive T cell response to viral infection and may shed light on the design of more effective vaccine strategies based on manipulation of γδT cells.

